# Density-Dependent Colour Scanning Electron Microscopy (DDC-SEM). Applications in the study of calcified tissues and visual impact

**DOI:** 10.1101/2024.09.16.613182

**Authors:** Elena Tsolaki, Luke Hunter, Adrian H Chester, Sergio Bertazzo

## Abstract

Scanning Electron Microscopy (SEM) is widely used as a technique for materials characterization. It has also been successfully applied to the imaging of biological samples, providing invaluable insights into the topography, morphology and composition of biological structures. A particular method combining different SEM detectors, named Density-Dependent Coloured SEM (DDC-SEM), has proven to be most useful for the identification and visualization of minerals in soft tissues. The method consists of a manipulation of original greyscale SEM images to produce coloured images that provide both topography and density information for samples with components of different densities. Here we provide a discussion on how to use DDC-SEM to aid the visualization and intuitive understanding of pathological calcification. This method has become popular not only for its scientific improvement of conventional SEM greyscale images, but also for its aesthetical merits.

**Lay summary:** Scanning Electron Microscopes (SEM) are widely used tools for examining biological materials such as human tissue. Like other electron microscopes, it only produces images in greyscale. SEM has been useful, for instance, in improving our understanding of calcific diseases. These diseases involve the build-up of mineral in the body’s soft tissues, and frequently affect the heart, kidneys, or eyes. This work provides a discussion on using a SEM technique known as Density-Dependent Colour Scanning Electron Microscopy (DDC-SEM), which enhances SEM images through scanning the same area with different detectors, assigning a unique colour to each detector’s output, and then overlaying these images.

## Introduction

The most recognizable calcified tissues on vertebrates are bone, teeth and osteoderms (in the form of turtle shells, for example)^1,2^. In addition to these natural calcified tissues, pathological calcification, or the formation of minerals in soft tissues, is a well-known process observed in the context of a number of diseases, including several cancer types, such as breast, ovary and prostate cancer^3-5^; Alzheimer’s disease^6^; tuberculosis^7^; chronic kidney disease^8,9^; aortic valve stenosis^10,11^; rheumatic heart disease^12^; among others^13^. One of the major challenges in the study of pathologic calcification is the identification or visualization of minerals in soft tissues^14-18^. This is due not only to size, but also to the fact that these minerals are not easily distinguished from the organic matrix^14-18^.

Scanning electron microscope (SEM) has been widely employed in the study of calcified tissues^14-23^, and a particular method combining different SEM detectors has proven to be most useful for the identification and visualization of minerals in soft tissues^14-18,21,22^. This method, named Density-Dependent Coloured SEM (DDC-SEM)^16^, consists of a manipulation of original greyscale SEM images to produce coloured images that provide both topography and density information (Fig. 1) ^16,24-^ 26.

**Figure 1.**
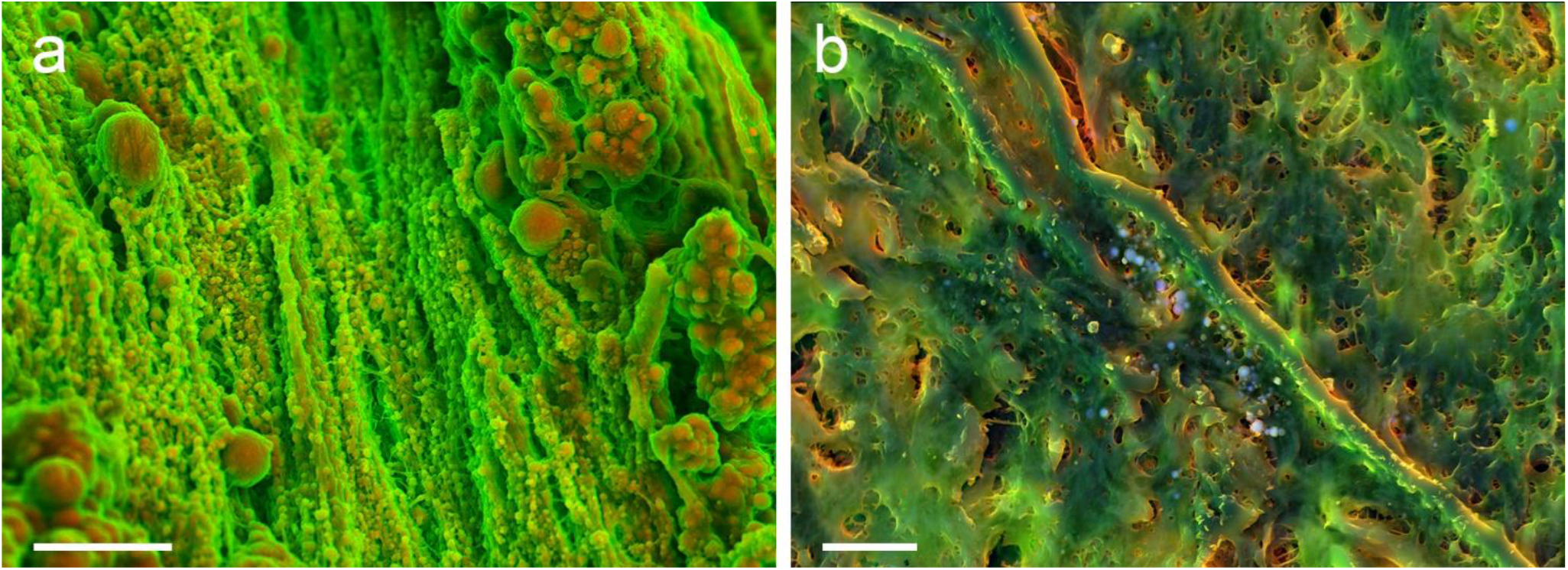
DDC-SEM micrographs of different samples, where organic and inorganic material can be distinguished based on their colour. (a) DDC-SEM of cardiovascular tissue, with orange being the inorganic, and green the organic components. Scale bar = 10 µm. (b) DDC-SEM of a histological slide of cardiovascular tissue, with green and red being the organic material and blue the inorganic material. Scale bar = 5 µm.

The approach leading to DDC-SEM was first suggested over a decade ago^25-27^ and was initially limited to randomly adding colour to greyscale SEM images merely for aesthetic purposes. Recently, its potential for the study of calcified tissue, and particularly for pathological calcification, has been rediscovered^14-18^. The DDC-SEM method allows for a clear visualization of organic and inorganic materials present in pathologically calcified soft tissues, owing to the contrast generated by the substantial differences in atomic number and density between the soft tissue and the mineral.

Here we discuss the sample preparation method and computational process to obtain DDC-SEM images from calcified soft tissues. The methodology to obtain DDC-SEM images is simple and can be applied using any SEM equipped with secondary electron (SE), and backscattered electron (BSE) detectors. We will use exemplars of pathological calcification samples, as the study of this disease has been producing images that capture the scientific value of DDC-SEM, as well as the aesthetical value of this method.

## Materials and Methods

The DDC-SEM method provides its best results when two materials of different composition and density are present. Indeed, calcified tissues (natural or pathological) present exactly these parameters, being formed from organic (cells and extracellular matrix) and inorganic (mineral) components.

Tissue samples were obtained by Old Brompton Hospital - London and from Oxford Heart Valve Bank at John Radcliffe Hospital – Oxford, the ethical approval for which was obtained for the original research paper^16^.

Biopsies of a few centimetres were randomly obtained from tissues, which were sectioned into small samples of two to five millimetres. After sectioning, samples were fixed in a 4% (mg/ml) formaldehyde in phosphate buffered saline solution at 4°C, for at least 24 hours. Samples were then dehydrated through a series of graded ethanol solutions; 20%, 30%, 40%, 50%, 60%, 70%, 80%, and 90% for a 10-minute interval each, followed by three changes of pure ethanol for 10 minutes each. After the dehydration procedure, samples were left to air dry. For pathological calcification studies, where the interest is on the composition and structure of the inorganic component, the visualization of the inorganic component is better if the samples are left to air dry (see Results and Discussion section). Dry samples were secured onto an aluminium stub using conductive carbon adhesive tape, carbon coated with 5 nm of carbon using a Quorum K975X thermal evaporator (Quorum Technologies, UK), and 5 nm chromium using a Quorum K575X sputter coater (Quorum Technologies, UK).

Samples were then imaged with a Zeiss LEO 1525 LG05B Field Emission Scanning Electron Microscope (Carl Zeiss, Germany) equipped with a secondary and a backscattered electron detector, and an in-lens detector. For DDC-SEM, samples should be imaged in a SEM equipped with secondary electron (SE), and backscattered electron (BSE) detectors. A voltage commonly used for imaging is 10 kV or higher. Whilst lower voltages are possible, the contrast of the BSE detector is, in general, not as pronounced. Micrographs were acquired routinely for the equipment in use and the images were superimposed using ImageJ^28^.

The same method may also be applied to histological slides of calcifying cell cultures.

## Results and Discussion

DDC-SEM images are formed from an overlay of two micrographs taken from the same region of a sample using different detectors. One micrograph should be acquired using the SE detector (or any variation of this, such as an in-lens detector, Fig. 2(a)) and another image should be acquired using a BSE detector (Fig. 2(b). A colour channel is assigned to the SE micrograph (in the example shown, the colour green is assigned to the in-lens image, Fig. 2(c)) and a distinct colour channel is assigned to the BSE micrograph (in this same case, red was assigned, Fig.2(d)). The superimposition of both colour images results in the inorganic material being presented in one colour (red/orange) and the organic material appearing in a different colour (green, Fig. 2(e)).

**Figure 2.**
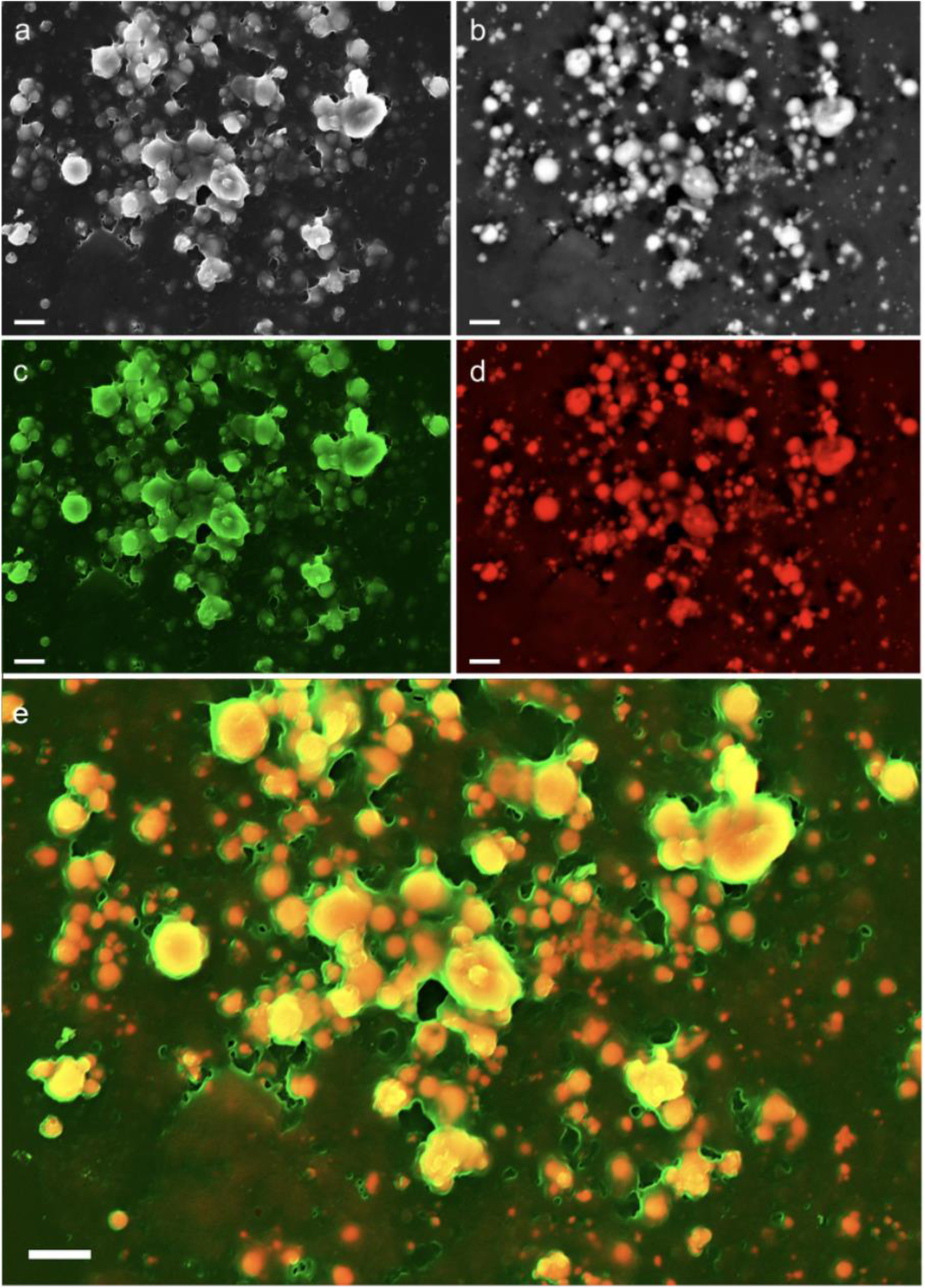
Density-dependent coloured electron micrographs of a mineralizing vascular smooth muscle cell culture model. (**a**) Original in-lens detector micrograph, providing topographic information. (**b**) BSE detector micrograph, showing dense material (minerals) in white, and organic material in black. (**c**) In-lens detector micrograph assigned the green channel. (**d**) BSE detector micrograph assigned the red channel. (**e**) Superimposed DDC-SEM image, with red and orange colours indicating the calcification and green indicating the organic matrix, whilst also providing topographic information. Scale bars = 2 µm.

DDC-SEM images are particularly useful when the micrographs that are superimposed had been acquired using in-lens/BSE (Fig. 2e). This is where the topographic information and the mineral present in the sample are quite clear (in this case, samples were air dried, on collapsed cells and extracellular matrix). This occurs because contrast levels of the BSE and in-lens images are considerably higher than the levels obtained between normal SE and BSE detectors. With a higher difference in contrast between the components of the sample, the final image DDC-SEM has richer colours.

One variation of the method that can add an interesting array of colours to the superimposed images is produced when multiple SEM detectors are available, generating multicolour DDC-SEM micrographs. For instance, green, red and blue channels may each be assigned to a different detector (in-lens, Fig. 3a, SE, Fig. 3b and backscattered, Fig. 3c). The resulting image (Fig. 3d) will present components in three colours.

**Figure 3.**
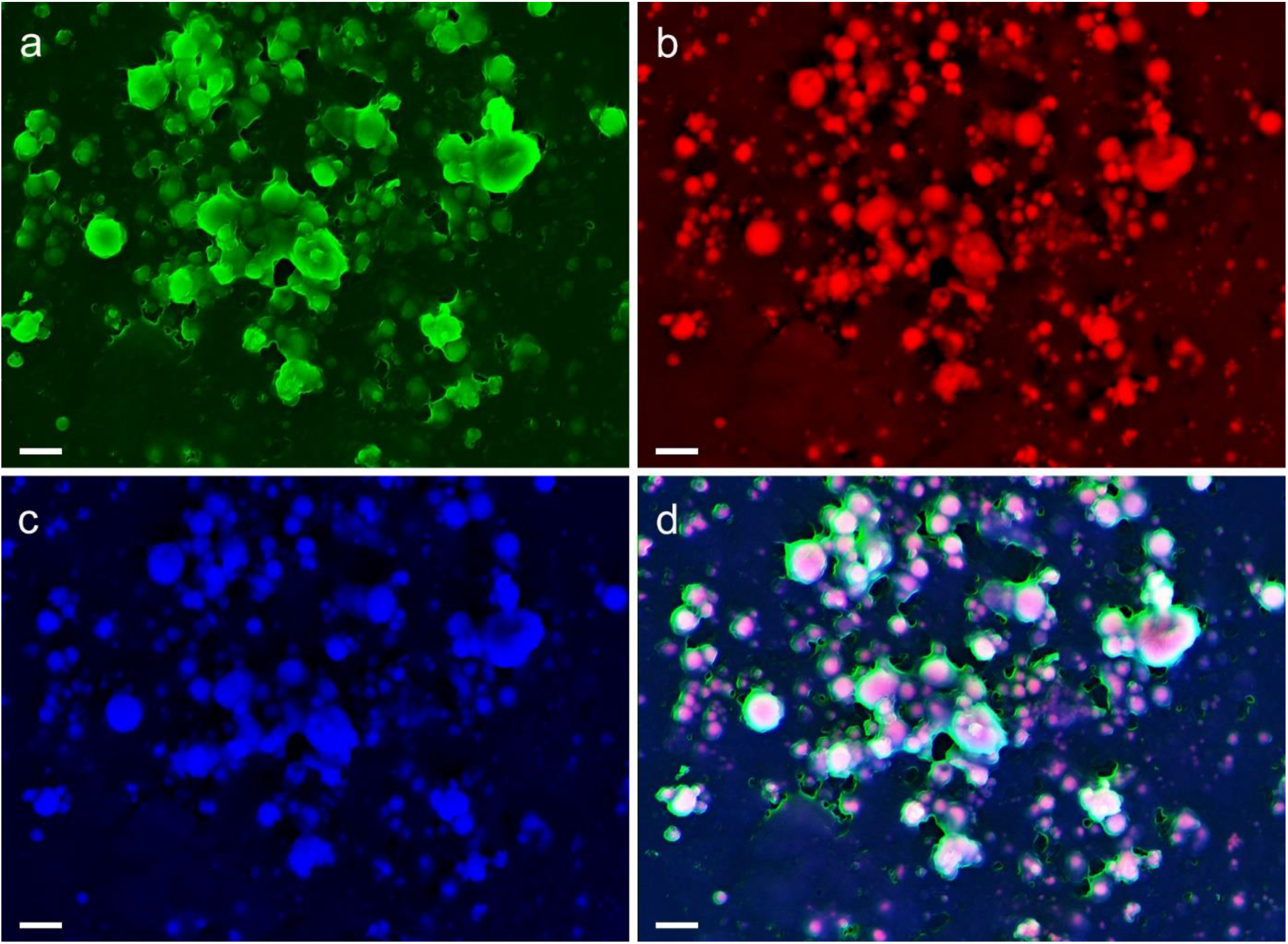
Density-dependent coloured electron micrographs of calcification in a mineralising vascular smooth muscle cell culture model. (**a**) In-lens detector micrograph assigned the green channel. (**b**) BSE detector micrograph assigned the red channel. (**c**) SE detector micrograph assigned the blue channel. (**d**) Resulting superimposed DDC-SEM image, where pink and purple colours indicate the calcification, while blue and green colours indicate the organic matrix. Scale bars = 2 µm.

Assigning colours to multiple modality-derived channels also provides an alternative to false-colour SEM images. In DDC-SEM, colours are obtained by the substitution of the signal (presented originally in greyscale) with the colour assigned, resulting in richer colours and higher contrast than the ones obtained with false colour. There are a few reasons why colours obtained on DDC-SEM are more vibrant when compared to false colour. First and mainly, this is because on false colour images colour is applied over the original greyscale image (instead of substituting the original image, as done in DDC-SEM). The overlay on a greyscale image fades the applied colour and reduces the contrast. Indeed, the assignment of colour directly to the signal obtained by each of the detectors keeps the original information about topography and density. Moreover, false colour images often rely on manual thresholding efforts and can highlight regions in an image as perceived by the person ascribing colours, allowing for bias and individual interpretation of structures and composition.

The colour coding on DDC-SEM images not only makes for an easier visualization of the different components of any sample presenting two components with distinct densities; DDC-SEM also helps to combine into a single image both the topographical information (obtained more clearly with an SE detector) and the density variation (obtained with a BSE detector). This variation of information can be illustrated by a line profile (Fig. 4a) along the same coordinates of the in-lens (Fig. 4aI), BSE (Fig. 4aII), and DDC-SEM images (Fig. 4aIII). The greyscale pixel values are plotted in Fig. 4b and show the feature differences as captured by the different detectors. Peaks in the BSE profile indicate the positions of particles which the in-lens profile will also detect if the topography is visible. However, the BSE profile does not clearly suggest topographical changes in the material, such as holes, and the in-lens detector fails to capture small particles just below the surface of the sample (Fig. 4b). DDC-SEM captures features from each and every one of these detectors. Hence, its profile indicates changes to both particles and topography in the sample. The grey value histograms from the in-lens and BSE detectors, when compared to the histograms from the DDC-SEM image, reveal that the latter integrates elements from both detectors. Indeed, its distribution predominantly falls between those of the in-lens and BSE histograms (Fig. 4c).

**Figure 4.**
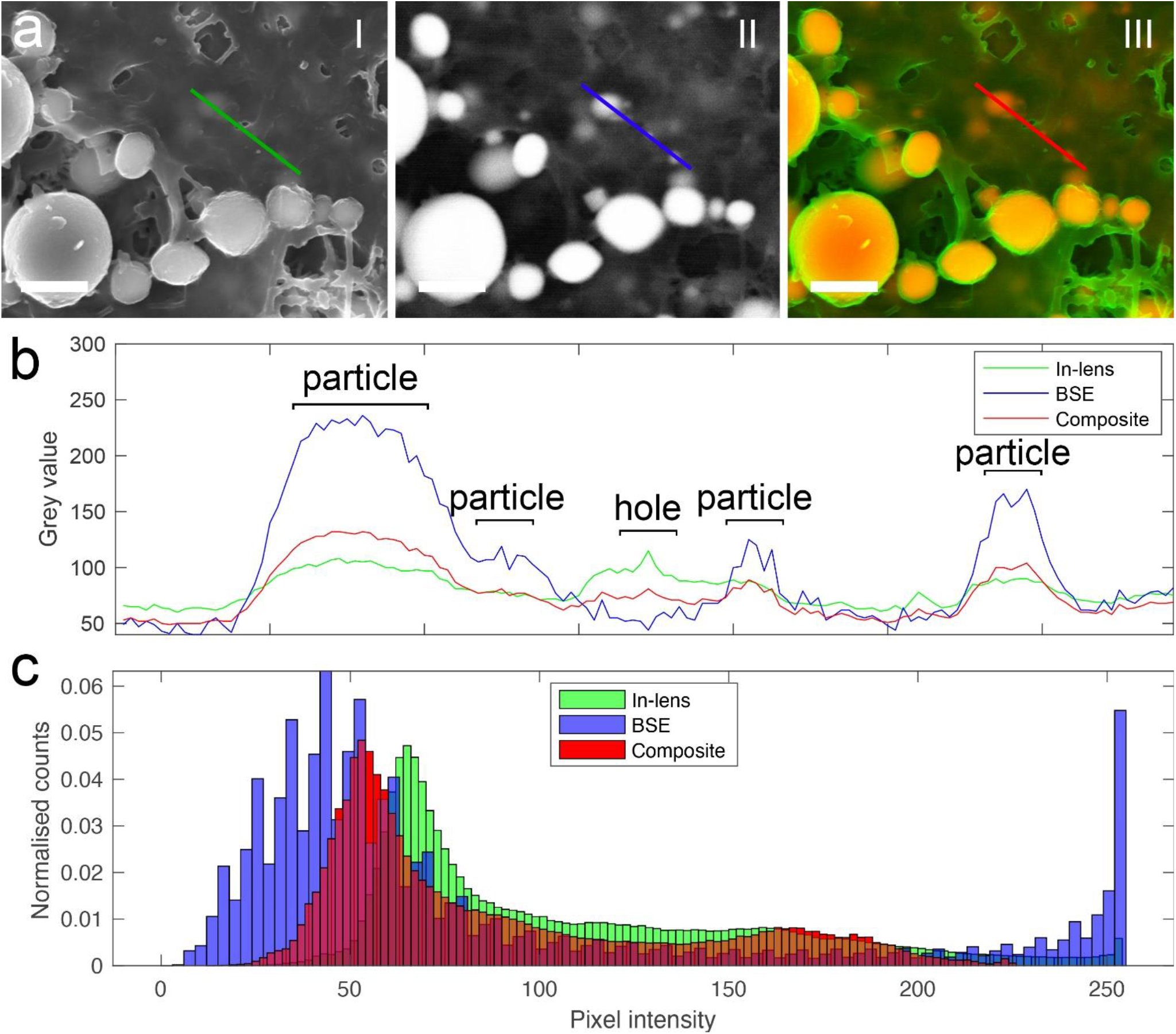
Electron micrographs of human aortic valve tissue and feature comparisons. **a** DDC-SEM micrograph produced from in-lens and BSE detectors. Line indicates position of line profile used. Scale bar = 1 µm. **b** Image profiles along line of composite DDC-SEM in Fig. 6a. **c** Histograms of image grey values obtained across entire visible image. Same images are compared in Fig. 6b.

To obtain high-quality DDC-SEM images, it is vital that samples present as little build-up of charge on their surface as possible (indeed, as required for any SEM micrography). The built-up charge not only distorts the image through the formation of horizontal lines and excessively bright areas (Fig. 5a), but also interferes in the colour assignment to the right location in the image (Fig. 5b).

**Figure 5.**
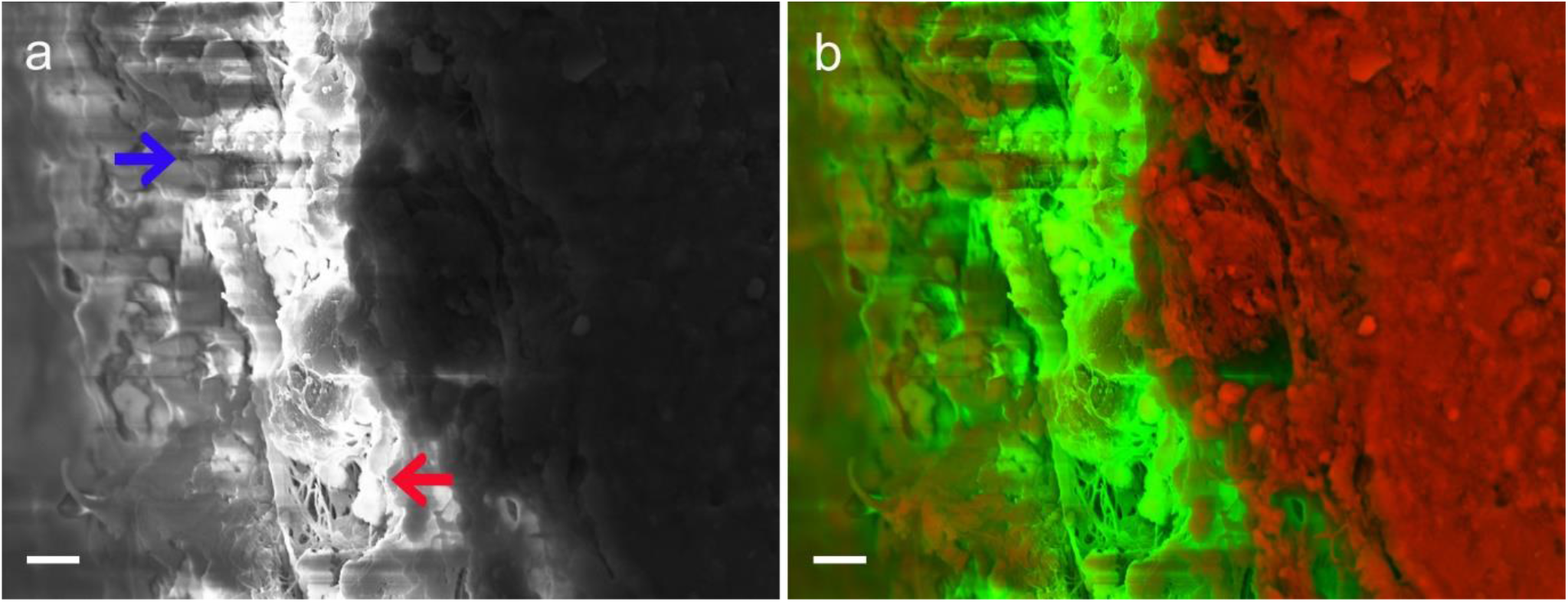
Images of inadequately prepared aortic valve tissue. **a** In-lens detector micrograph of a sample where no silver paint was applied, showing the ‘charging’ effects on the sample: horizontal lines (blue arrow) and excessively bright regions (red arrow). **b** Poor quality DDC-SEM micrograph of the same sample, where charging effect is clearly observed. Scale bars = 4 µm.

Adequate dehydration and drying methods must also be considered for better-quality DDC-SEM micrographs. The most common approaches for dehydration are critical point drying and hexamethyldisilazane (HMDS) dehydration^29,30^. These methods have been developed to maintain, as much as possible, the original structures from biological samples as they would be found in the hydrated state. While critical point dry would not affect the density of the overall sample, HMDS can have a considerable detrimental effect on the contrast obtained by the BSE detector.

In the study of pathological calcification, it is often desirable to obtain most information about the inorganic component of a sample. Air drying a biological sample that contains minerals will cause the organic material to shrink and collapse around the mineral, distorting the information about the organic component on the sample (cells and extracellular matrix), and on the other hand enhancing the amount and quality of information about the mineral. Therefore, air-dried samples can provide DDC-SEM of calcified samples where the inorganic material can be easily visualized and with increased details.

A comparison between air-dried samples and HMDS dried samples shows that the latter allows for visualization of greater detail of the organic component of the sample, as expected, but makes it hard to distinguish whether the structures observed are organic or inorganic (Fig. 6a and Fig. 6c). Indeed, practically only the minerals on the surface of the sample can be identified (Fig. 6c and Fig. 6e).

**Figure 6.**
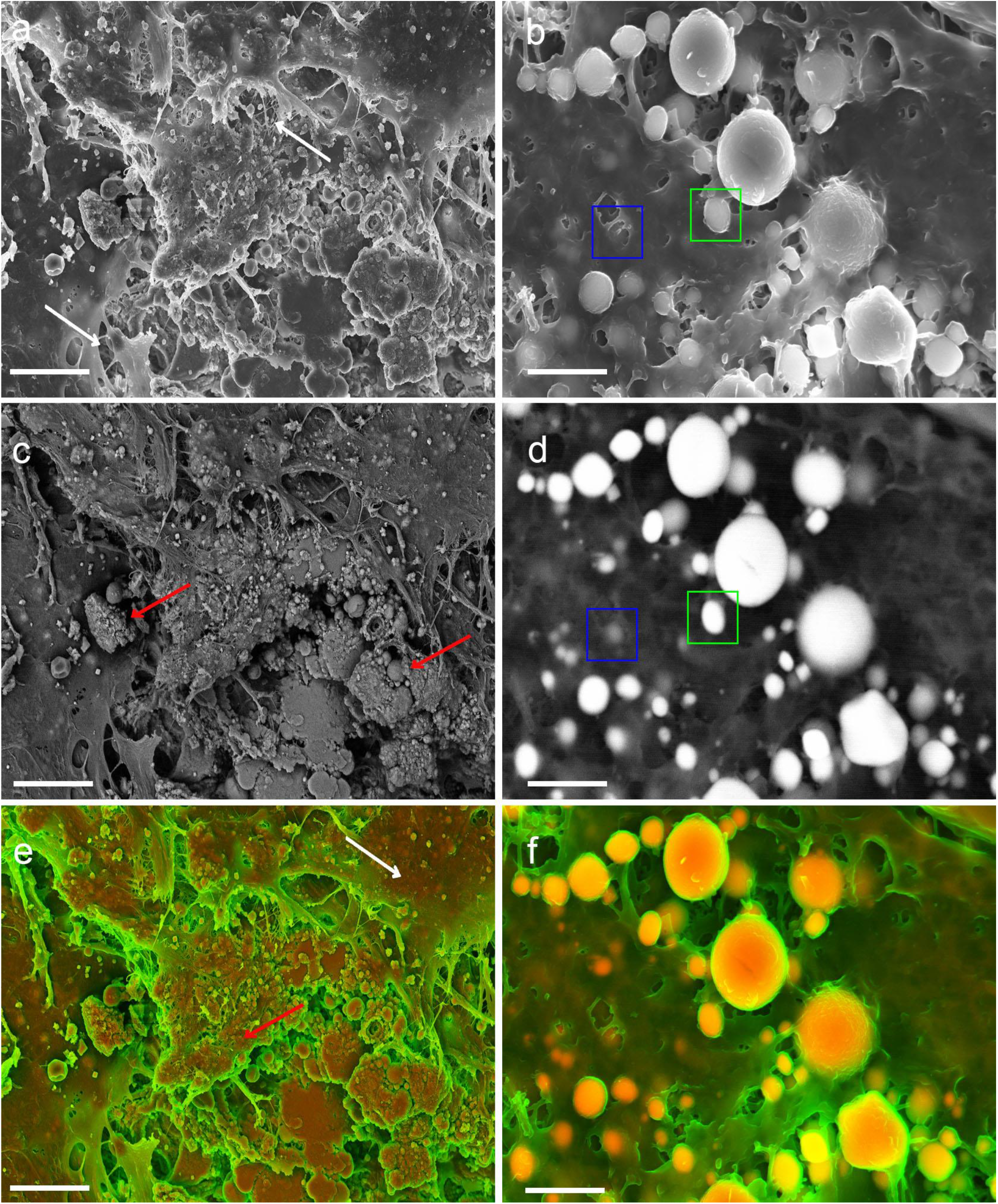
Electron micrographs of human aortic valve tissue. **a** In-lens detector micrograph of a HMDS treated sample, where the organic microstructure can be visualised (white arrows) Scale bar =4 µm. **b** In-lens detector micrograph of an air-dried sample, where the organic material has collapsed, allowing for calcification to be more easily visualised. Scale bar = 2 µm. **c** BSE micrograph of the area shown in **a**, where some calcification can be observed on the surface of the sample (red arrows). Scale bar = 4 µm. **d** BSE micrograph of the area shown in **b**, where calcification can be easily visualised with marked contrast differences between the organic and inorganic material. Calcifications that were either closer to the surface (green square) or deeper in the tissue can be equally well identified (blue square). Scale bar= 2 µm. **e** DDC-SEM micrograph produced by assigning the green channel to the image in **a** and the red channel to the image in **c**, with the poor contrast of the BSE micrograph resulting in an image where it is hard to distinguish between the organic (white arrow) and inorganic (red arrow) material based on colour. Scale bar = 4 µm. **f** DDC-SEM micrograph produced by assigning the green channel to the image in **b** and the red channel to the image in **d**, where the organic and inorganic materials can be easily distinguished based on colour. Scale bar = 2 µm.

As expected for the air-dried sample, the organic matrix collapses (Fig. 6b), allowing for minerals to be easily visualized (Fig. 6b), even if located under organic material (Fig. 6d). A comparison between DDC-SEM micrographs produced for HMDS prepared samples (Fig. 6e) and the air-dried samples (Fig. 6f) reveals how the latter allows for easy identification of the morphology and location of calcified deposits.

## Conclusion

The use of DDC-SEM can assist in the study of pathological calcification and contribute towards advances in that field, as it allows for a clear and prompt visualization of minerals present in biological tissue samples. Indeed, this method discussed here can be applied to any type of sample that has components with a considerable difference in density. This imaging method is already being increasingly adopted for the study of pathological calcification and in biomineralization systems. It has become a popular method not only for its scientific improvement of conventional SEM greyscale images, but also for its aesthetical merits, as demonstrated by the large number of journal covers recently featuring DDC-SEM images, and frequent use of these images on scientific and academics websites. Finally, it is important to highlight that the DDC-SEM method can be incorporated into the visualization and characterization of minerals alongside other methods, contributing to a holistic view of mineralized tissues at large.

